# Barley C2-Domain Abscisic Acid-Related protein CARa supports susceptibility to *Blumeria hordei* and localizes to the extrahaustorial membrane

**DOI:** 10.64898/2025.12.04.692366

**Authors:** Mariem Bradai, Christopher McCollum, Ralph Hückelhoven

## Abstract

Rho GTPases are key regulators of cellular signalling processes in eukaryotes. In plants, Rho of plants (ROP) proteins function in cell polarization, hormone signalling, and plant immunity or susceptibility to diseases. The barley (*Hordeum vulgare*) ROP protein RACB is involved in susceptibility towards the biotrophic ascomycete fungus *Blumeria hordei* (*Bh*) and supports the accommodation of fungal haustoria. In healthy barley, RACB plays a role in cell development, a function that the pathogen may co-opt for cellular ingrowth of its haustorium into intact epidermal cells of barley. The majority of identified functions of GTP-bound activated RACB appear to be mediated by scaffold proteins. One of them, the ROP Interactive Partner b (RIPb, synonym: Interactor of Constitutive Active ROP b, ICRb) binds to RACB-GTP and co-localizes with RACB at the site of fungal host cell entry. RIPb interacts with RACB-GTP via its C-terminal coiled-coil CC2 domain (RIPbCC2) at the plasma membrane. Here, we show that a barley Calcium-Dependent Phospholipid Binding 2 (C2)-Domain Abscisic Acid-Related protein (CARa) interacts with RIPb and its RIPbCC2 domain in *planta*. Transient knockdown of CARa renders barley epidermal cells less susceptible to invasion by *Bh*, whereas overexpression supports fungal haustorium accomodation. Upon fungal attack, CARa is recruited to the site of fungal attack and localizes around the haustorial neck and at the extrahaustorial membrane. These findings further support that *Bh* profits from host ROP signalling components and CARa. CARa might connect RACB-RIPb-signalling to membrane organization for support of fungal infection structures in barley epidermal cells.

## Introduction

During infection, the powdery mildew fungus *Blumeria hordei* (*Bh*), which is an obligate biotrophic ascomycete parasite, grows on the areal plant organs of cultivated barley (*Hordeum vulgre*) (Troch et al., 2014). During the asexual life cycle of *Bh*, conidia land on a leaf surface and germinate with a primary non-infectious and a second appressorial germ tube, from which the fungus attempts to penetrate. If the attempt is successful, the fungus penetrates the plant cell wall and establishes a specialized invasive structure known as haustorium (Glawe, 2008). The latter serves as feeding structure for the fungus during all stages of pathogenesis and for the release of virulence effectors (Catanzariti et al., 2007). Haustoria are surrounded by an extrahaustorial matrix and an extrahaustorial membrane (EHM), which forms a continuum with the host’s plasma membrane (PM) but both membranes are structurally and compositionally distinct (Micali et al., 2011; O’Connell & Panstruga, 2006). In some plant-powdery mildew pathosystems, including *Arabidopsis thaliana* (hereafter referred to as Arabidopsis) interactions with *Golovinomyces orontii* (*Go*), and *Erysiphe cichoracearum* (*Eo*), typical PM markers are excluded from the EHM, despite their presence in adjacent PM (Koh et al., 2005; Micali et al., 2011). In *Bh* pathogenesis, the EHM appears to be plant-derived and is closely associated with the host endoplasmic reticulum (ER), sharing multiple features with it (Micali et al., 2011; Kwaaitaal et al., 2017). Interestingly, the EHM withstands chemical treatments that normally disrupt the PM and is devoid of all examined proteins present in the PM (Gil & Gay, 1977; Koh et al., 2005; Micali et al., 2011). Diverse models have been proposed for the EHM biogenesis; invagination of the plant PM followed by membrane differentiation, or de novo formation via targeted trafficking of vesicles or lipids of diverse origin (Koh et al., 2005; Kim et al., 2014; Inada et al., 2016; Berkey et al., 2017; Kwaaitaal et al., 2017). Despite their importance in the plant-pathogen interaction, the origin and nature of these extrahaustorial membranes remain controversial. To date, only a limited number of plant proteins have been shown to localize to the EHM associated with powdery mildew fungi. The Arabidopsis resistance to powdery mildew 8.2 protein (RPW8.2), an R-protein that provides broad-spectrum resistance to *G. orontii*, is found to be transiently targeted by the SNARE protein VAMP721/722-decorated vesicles within the cell to the EHM (Wang et al., 2009; Kim et al., 2014). Additionally, and in the same Arabidopsis-powdery mildew system, ARA6, a plant-specific RAB5 GTPases (RAB5, RABF) is localized to the EHM (Inada et al., 2016). Specific RAB GTPases regulate individual trafficking pathways, with RAB5 as a key regulator of diverse endocytic processes (Ebine & Ueda, 2009). A similar RAB5/RABF GTPase but also SAR1 and RABD2a GTPases involved in endoplasmic reticulum membrane trafficking localize also to the EHM of *Bh* in barley (Inada et al. 2016; Kwaaitaal et al. 2017).

Plant monomeric RHO GTPases (rat sarcoma homologues, also called RAC for rat sarcoma (RAS)-related C3 botulinum toxin substrate) are small monomeric G-proteins that form a unique subfamily of Rho in plants (ROP). They are involved in cell polarity, vesicle trafficking, cytoskeleton organization and plant immunity (Engelhardt et al., 2020). G-proteins can switch between an inactive, GDP-bound state and an activated GTP-bound conformation (Berken & Wittinghofer, 2008). If activated, ROP proteins interact with downstream effectors (also called executors to distinguish from pathogen effectors) and regulate a multitude of cellular mechanisms in plants (Engelhardt et al., 2020; Feiguelman et al., 2018). In barley, the overexpression of a constitutively activated ROP protein (CA RACB) enhances the penetration success of *Bh* into epidermal barley cells, whereas RNA interference mediated silencing of *RACB* reduces penetration success and haustorial size. This suggests that RACB functions as a host susceptibility factor in the powdery mildew pathosystem (Hoefle et al., 2011; Schultheiss et al., 2002, 2003). Beyond its role in susceptibility, RACB also regulates the development of stomatal subsidiary cells and root hair outgrowth (Hoefle et al., 2011; Scheler et al., 2016). Both processes require single-cell polarization, suggesting that RACB-mediated susceptibility to *Bh* is linked to its function in cell polarity. However, RACB does not act alone in this process; scaffold proteins likely facilitate its interaction with downstream executors (ROP effectors) during fungal haustoria accommodation (Engelhardt et al., 2020). In barley, eight RICs (ROP-Interactive and CRIB-(Cdc42/Rac Interactive Binding) motif containing) proteins are identified and at least two of them interact with RACB (Engelhardt et al., 2023; Schultheiss et al., 2008). Interestingly, RIC171 and CA RACB act similarly but not additively on *Bh* penetration success, and both form a complex at the site of fungal attack, suggesting their function in the same signaling pathway (Hückelhoven & Panstruga, 2011; Schultheiss et al., 2008). Similarly, RIC157 and CA RACB support fungal invasion and co-localize at the haustorial neck (Engelhardt et al., 2023). ICR/RIPs (Interactor of Constitutive Active ROP/ROP Interactive Partners) form another class of scaffold proteins (Lavy et al., 2007; S. Li et al., 2008), and barley RIPb apparently supports RACB signaling during haustorium accommodation (McCollum et al., 2020). In Arabidopsis, ICR/RIPs are required for cell polarity in ROP signaling to control polarized pollen tube growth and share a carboxy-terminal coiled-coil CC2 domain with a QWRKAA amino acid motif described as involved in the ROP-RIP interaction (Lavy et al., 2007; S. Li et al., 2008). RIPb, one of three barley ICR/RIP scaffold protein members, also contains a conserved CC2 domain and a QWRKAA motif. Both full length RIPb and a truncated form harboring the CC2 domain (RIPbCC2) interact with RACB and were recruited to the cell periphery by plasma membrane-attached CA RACB (McCollum et al., 2020). Similarly to RIC157 and RIC171, RIPb co-localizes with RACB at the site of fungal attack and its transient overexpression in epidermal cells enhances susceptibility to *Bh* (McCollum et al., 2020). In Arabidopsis, a positively charged amino acid motif at the C-terminus is important for membrane localization of AtRIP1/ICR1 and regulation of polarized pollen tubes tip growth (S. Li et al., 2008). Interestingly, RACB possess also a carboxy-terminal positively charged polybasic region (PBR). This feature is conserved among other plant ROPs and responsible for anionic phospholipid interaction (Schultheiss et al., 2003); (Weiß et al., 2025); (Platre et al., 2019). RACB indeed binds to anionic phospholipids and RACB-5Q, in which five charged lysine residues in the PBR were substituted with glutamine residues fails to interact with the anionic phospholipids. Additionally, the CA RACB-5Q can no longer localize at the cell periphery and is not able to support susceptibility towards *Bh* as polybasic CA RACB (Weiß et al., 2025). Together, these findings highlight the importance of PM-localization for the function RACB as a susceptibility factor.

Here, we show that RIPb and its RIPbCC2 domain interact with a barley Calcium-Dependent Phospholipid Binding 2 (C2)-Domain Abscisic Acid-Related protein (CARa) *in planta*. CARa binds to phosphatidylserine *in vitro*. Strikingly, CARa labels the EHM, and we found that is a susceptibility factor with potential function in the RACB-RIPb pathway. We discuss potential modes of action in which RACB, RIPb, and CARa meet in phospholipid-rich zones at the PM to promote *Bh* during host cell penetration.

## Results

### Identification of CAR Proteins in Barley

ICRs/RIPs are described as a scaffold proteins and suggested to connect activated ROPs with downstream ROP effectors (Lavy et al., 2007; McCollum et al., 2020). To further understand the function of the RACB-RIPb connection, we performed a yeast two-hybrid screening against a barley cDNA library using barley RIPb as a bait. Among the obtained interactors, we identified a C2-domain ABA (abscisic acid)-related protein (CAR). In Arabidopsis, all 11 CAR proteins identified contain a single predicted lipid-binding C2 domain (Rodriguez et al., 2014). Based on this, we performed an *in silico* analysis to identify further barley CAR family members, and compare them to Arabidopsis. After BLAST searches with protein sequence of the identified barley CAR (HORVU.MOREX.r3.5HG0469960), we identified 45 hits with C2 domains in barley (reference: Morex v3 in EnsemblPlants). Six of them show similarity to Arabidopsis CARs, harbour only a single C2 domain and show no further conserved protein domains. We therefore considered them canonical CAR proteins (Figure 1a). Based on prediction using InterPro (Blum et al., 2025) we found that the six C2 domain proteins show similarity to the 11 CAR family members of Arabidopsis. Because we found only six CARs in barley but 11 in Arabidopsis, and their phylogenetic relation is not unambiguous (see below), we named the barley CARs *Hv*CARa to *Hv*CARf instead of numbering them from one to six (supplementary Table S1). We also identified six CAR proteins in *Brachypodium distachyon* and seven CAR proteins in rice (*Oryza sativa*) (supplementary Table S1). The alignment of the amino acid sequences of CARs from barley, Brachypodium, and rice shows that they possess a conserved predicted C2 domain and no further protein domains. They contain conserved aspartate residues common to the C2 domain of protein kinase C and previously described Arabidopsis CAR proteins (Figure 1a; supplementary Figure S1). These residues were shown to be important for Ca^2+^ ion binding and phospholipid interaction. Furthermore, both grass and Arabidopsis proteins possess conserved aromatic and lysine residues that are responsible for phospholipid binding specificity (Figure 1a) (Rodriguez et al., 2014; Verdaguer et al., 1999; Yung et al., 2015). However, the exact positions of lysine residues were not strongly conserved between *Hv*CARa and the CAR proteins, *Os*GAP1/OsCARe and *At*CAR4, for which crystal protein structures have been described (Rodriguez et al., 2014; Yung et al., 2015) (Figure S1). Pairwise comparisons show that the CAR protein sequence identity within each cluster ranged from 51% to 87% (Percent Identity Matrix, Clustal Omega). By contrast, protein identity between clades was comparatively lower. This pattern supports the subgroup clustering observed in the phylogenetic analysis (Figure 1b, supplementary Table S2).

**Figure 1:**
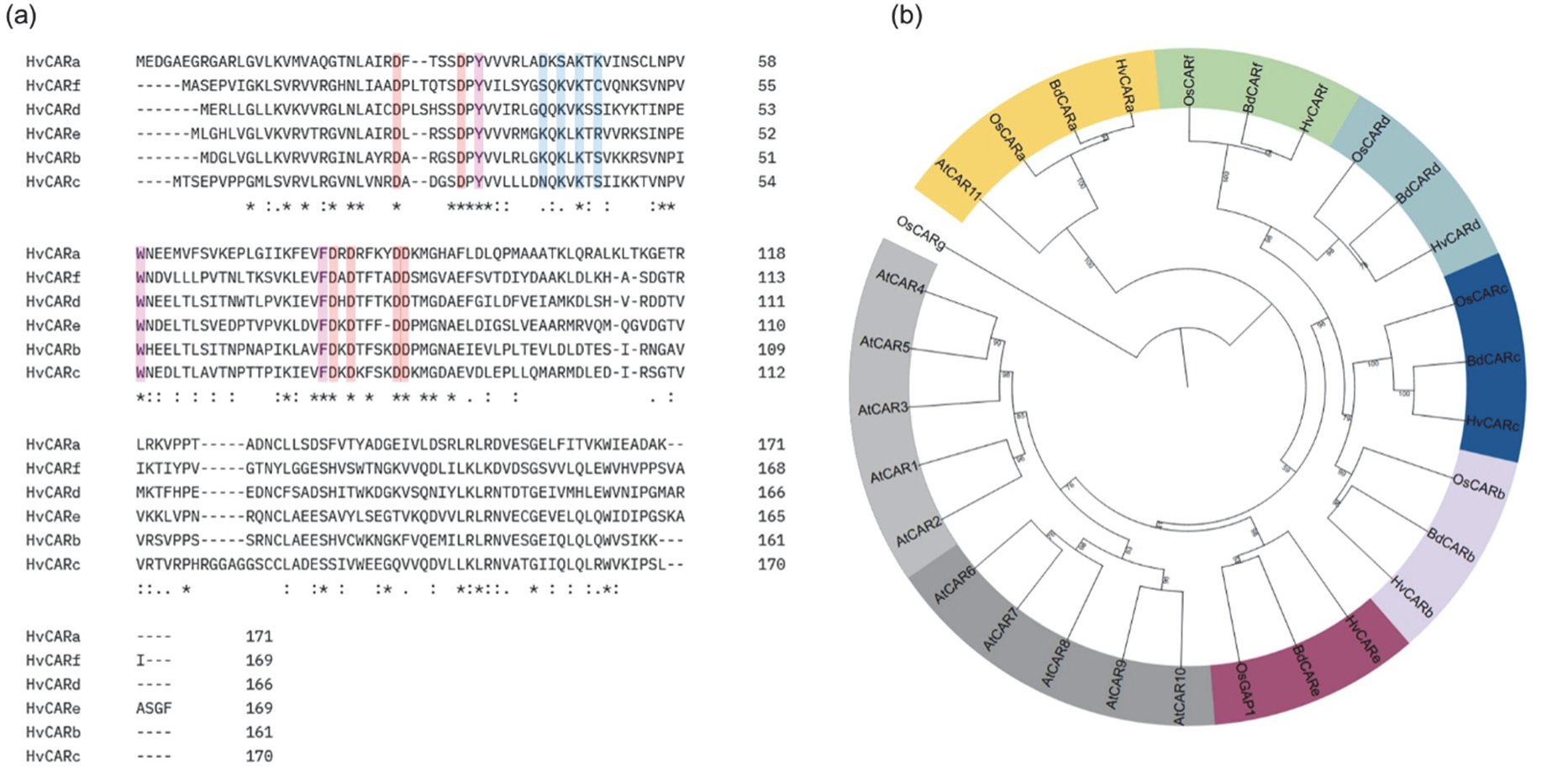
Phylogenetic and sequence analysis of barley CAR proteins. (a) Multiple sequence alignment (MSA) of *Hordeum vulgare* (Hv) CAR proteins generated using MUSCLE (Clustal EMBL format). Aspartate residues involved in Ca²⁺ binding and phospholipid interaction are highlighted in red, aromatic and lysine residues responsible for phospholipid binding specificity are highlighted in purple and sky blue, respectively. Protein identifiers correspond to EnsemblPlants accessions, and residue annotation is based on alignment with *Arabidopsis thaliana* CAR proteins (see Figure S1). (b)Phylogenetic tree based on CAR protein sequences from *Hordeum vulgare* (Hv), *Arabidopsis thaliana* (At), *Oryza sativa* (Os), and *Brachypodium distachyon* (Bd). Protein sequences were were retrieved from EnsemblPlants and TAIR databases and aligned using MAFFT (Katoh & Standley, 2013), and the phylogenetic tree was inferred using IQ-TREE (Nguyen et al., 2015) with the best-fit substitution model automatically selected. The resulting tree was visualized using iTOL (Letunic & Bork, 2021).

### CARa is a bona-fide lipid binding protein in the barley leaf epidermis

To gain insight into the structural organization of barley CARa, we generated a three-dimensional protein model using AlphaFold (Jumper et al., 2021) and visualized it with ChimeraX (Pettersen et al., 2021) (Figure 2a). The predicted structure shows all characteristic features of a C2 domain, including a compact bipartite β-sandwich scaffold composed of two four-stranded β-sheets arranged in the β4-1-8-7 and β3-2-5-6 topology (topology II), as previously described for Arabidopsis AtCAR4 or rice OsGAP1/OsCARe (Rodriguez et al., 2014; Yung et al., 2015; Diaz et al., 2016). At one end of the protein, three surface-exposed loops form cavities that coordinate two calcium ions, as observed in CAR4 and CAR6 (Diaz et al., 2016; Khan et al., 2019). This region also contains some cationic lipid-binding amino acids and four conserved aspartate residues that constitute two calcium-binding pockets. One of these sites has high calcium affinity and serves a structural role, while the second site binds calcium with lower affinity (Rodriguez et al., 2014; Yung et al., 2015). Together, these features confirm that barley CARa likely adopts a typical C2-domain architecture, supporting its functional similarity to other plant CAR proteins. Additionally, the alignment of amino acid sequence of CARs from Arabidopsis and grasses shows a conservation among the residues responsible for lipid interaction and Ca^2+^ ions binding (Figure S1), which suggests that the barley CARa might interact with phospholipids. To determine whether CARa binds to phospholipids, a recombinant GST-CARa protein was expressed in *E*. *coli*, and purified via glutathione columns. We performed *in vitro* lipid-binding assays as previously described for barley RACB and RACB-binding proteins (Weiß et al., 2025), except that the blocking solution used contained bovine serum albumin (BSA). The result of the lipid-binding assay shows that barley CARa binds to phosphatidylserine (PS) similarly to the CAR4 of Arabidopsis (Rodriguez et al., 2014) (Figure 2b).

**Figure 2:**
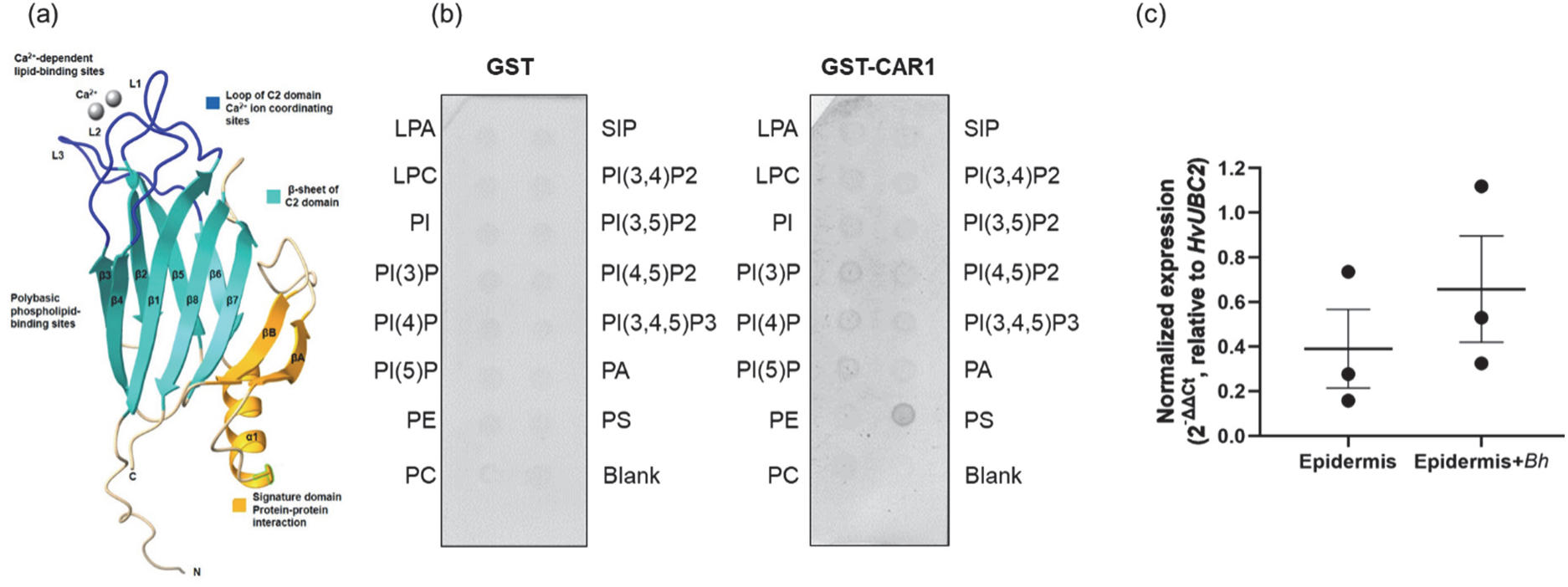
CARa possess all the conserved domain of a typical C2-domain and binds to phospholipids in vitro.

To explore the role of barley *Hv*CARs, gene expression patterns across various tissues was examined *in silico* using RNA sequencing (RNA-Seq) data from the barleyGenes database provided by the James Hutton Institute (*barleyGenes - Barley RNA-Seq Database*, n.d.; Rapazote-Flores et al., 2019). The data show that most of the *HvCARs* are broadly expressed with partially elevated fragments counts in various plant organs or tissues (supplementary Table S3). Given that the leaf epidermis of barley represents a critical interface for powdery mildew interaction, we focused on the expression of *HvCARs* in this tissue and found that *Hv*CARa, *Hv*CARd, *Hv*CARe, and *Hv*CARf are expressed in the epidermis of barley leaves (supplementary Table S3). In order to examine the gene expression of *HvCARa* during interaction with *Bh*, we conducted a reverse transcription-quantitative PCR (RT-qPCR) on transcripts from peeled leaf epidermis from 10-day-old seedlings of the barley cultivar Golden Promise, either non-inoculated or inoculated with conidia of *Bh.* RT-qPCR results show that *HvCARa* showed slight but not significant increase of *HvCARa* transcript abundance in inoculated epidermis compared to the non-inoculated epidermis (Figure 2c).

### CARa interacts with RIPb and its RIPbCC2 domain *in planta*

A yeast two-hybrid screening initially identified barley CARa as a potential interactor of RIPb. However, when we attempted to revalidate the interaction in a more stringent targeted Y2H assay, we did not observe yeast growth (Figure S2). This discrepancy suggests that the interaction may depend on specific cellular conditions or modifications that are absent in the yeast system or unidentified additional host factors co-expressed in the original screening. As an additional way to test for protein-protein interaction *in planta*, we performed bimolecular fluorescence complementation (BiFC) assays in which we tested cYFP-CARa against nYFP-RIPb (Figure 3a). This way, we obtained YFP fluorescence restoration. When we used dominant negative RACB, DNRACB, as negative control neither RIPb nor CARa lead to BiFC, whereas the positive control, constitutively activated RACB (CARACB) together with RIPb complemented YFP fluorescence (Figure 3a). We additionally tested if the C-terminal part of RIPb, RIPbCC2, interacts with CARa. In this experiment, YFP fluorescence was also restored under co-expression of RIPbCC2 and CARa (Figure 3b). These findings suggest a direct interaction between CARa and RIPb as well with the RIPbCC2 coiled coil protein domain *in planta.* Interestingly, the BiFC signal from CARa and full length RIPb was partially visible at the cell periphery and at microtubules. In contrast, CARa-RIPbCC2 complex was restricted to the cell periphery and to some weaker cytoplasmic signals (Figure 3a/b). This was similar to what was reported earlier for CARACB-RIPb and CARACB-RIPbCC2 complexes (McCollum et al., 2020) and confirmed here in our positive controls. Additional ratiometric quantification of the BiFC signal versus the co-expressed mCherry (mCh) in at least 25 cells showed a strong and significant difference between cells where RIPb or RIPbCC2 was co-expressed with CARa, and cells where RIPb/RIPbCC2 was co-expressed with DNRACB (Figure 3c and 3d). To independently confirm this interaction, we conducted Förster-Resonance Energy Transfer Fluorescence Lifetime Imaging Microscopy (FRET-FLIM) in barley. We transiently expressed meGFP-RIPb as donor and co-expressed mCh-CARa as acceptor. FLIM was measured at the cell periphery. Excitation of the donor only without any acceptor present resulted in a GFP fluorescence lifetime of about 2.6 ns on average (Figure 3e). As a negative control for interaction with RIPb, we included mCh-MAGAP1, a RACB interactor that partially co-localizes but does not interact with RIPb (Hoefle et al., 2011; McCollum et al., 2020). This resulted in slight reduction of GFP fluorescence lifetime. However, co-expression of mCh-CARa significantly and more strongly reduced the fluorescence lifetime of meGFP-RIPb, confirming a direct protein-protein interaction *in planta* (Figure 3e). We also tested by FRET-FLIM if the CC2 domain of RIPb interacts with CARa. In those experiments, mCh-CARa also reduced the lifetime of meGFP-RIPbCC2 when compared to the negative control, C-terminally mCherry-tagged glutathione S-transferase (mCh-GST) (Figure 3f).

**Figure 3:**
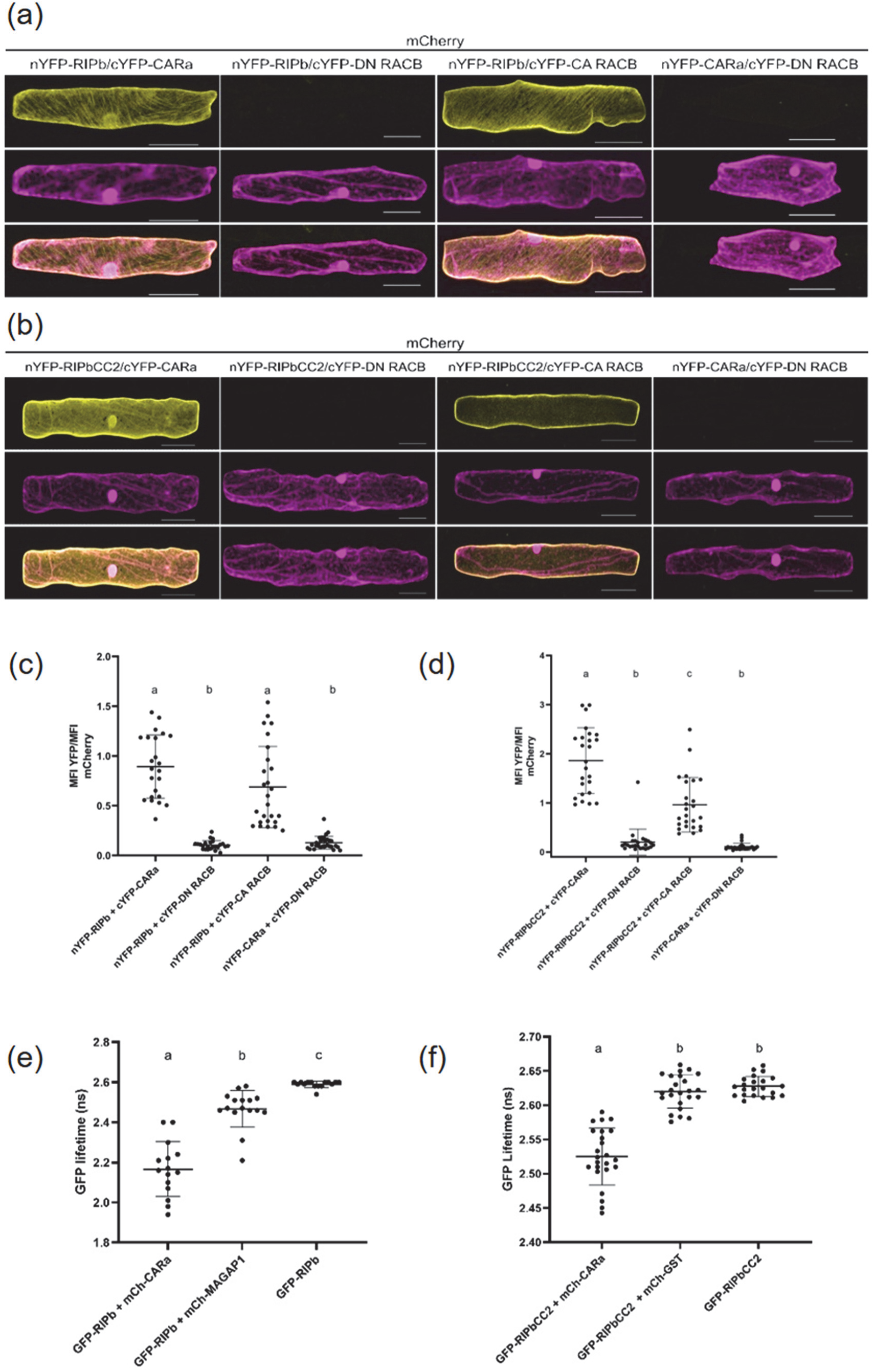
CARa protein interacts with RIPb, RIPbCC2 *in planta*. (a,b) Interaction of RIPb and RIPbCC2 with CARa in BiFC assays. Single epidermal cells were transiently transformed by particle bombardment with split-YFP constructs in the indicated combinations. Images represent typical cell recordings of minimum 20 cells per experiment and from two independent transformation experiments with similar results. Images represent z-stacks of 25 confocal sections. Image brightness was uniformly enhanced post-scanning for better visibility. Scale bar = 50µm. (c,d) Quantification of BiFC signals from images were taken with constant settings. Signal intensity (mean fluorescence intensity [MFI]) was measured over a region of interest at the cell periphery. The ratio between split-YFP and free mCherry signal was calculated. Signals were measured in minimum 20 cells for each construct. Letters indicate significance by one-way ANOVA (Tukey’s multiple comparison test; P < 0.01). (e,f) Interaction of GFP-RIPb and GFP-RIPbCC2 with mCherry-CARa as measured in FRET-FLIM experiments. Barley epidermal cells were transiently transformed via particle bombardment with expression constructs for either GFP-RIPb or GFP-RIPbCC2 as FRET-donors and mCherry-CARa as a FRET acceptor. mCherry-MAGAP1 or GST-mCherry served as no-interaction controls. FRET-FLIM measurements were conducted at the cell periphery of the equatorial plane of barley epidermal cells 1 day after transformation. Letters indicate significance by one-way ANOVA (Tukey’s multiple comparison test; P < 0.01). Measured cells were collected in at least three independent biological replicates.

### *Hv*CARa acts as a susceptibility factor towards *Bh*

In order to investigate the involvement of CARa in interaction with the fungal parasite *Bh*, we transiently overexpressed CARa in barley leaf epidermal cells, followed by inoculation with *Bh* conidia, fixation and staining for GUS activity of transformed cells. Fungal penetration success was rated by presence of a haustorium in a GUS-expressing cell. Unsuccessful attacks were counted when the fungus built an appressorium on a GUS-expressing cell but no haustorium was formed by 48 h post inoculation. Data showed a significant increase of the fungal penetration success in CARa overexpressing cells when compared empty vector controls. The relative fungal success increased by 60% in CARa overexpressing cells when compared to the empty vector controls (Figure 4a). We then created RNAi-mediated gene silencing construct for *CARa*, and we assessed the efficacy of the silencing by measuring the reduction of GFP fluorescence intensity of GFP-CARa. Barley epidermal cells were co-transformed with GFP-CARa and the RNAi silencing construct CARa-RNAi. Co-delivery of the CARa-RNAi vector led to more than 90% reduction of GFP fluorescence compared to the control vector (supplementary Figure S3). Silencing of *CARa* by RNAi and subsequent inoculation showed an effect opposite to the overexpression. Silencing *CARa* decreased the relative fungal success rate in penetration and haustorium formation by about 30% (Figure 4b). Thus, both single cell silencing and overexpression of CARa influence fungal infection success suggesting that CARa supports accommodation of fungal haustoria in barley epidermis.

**Figure 4:**
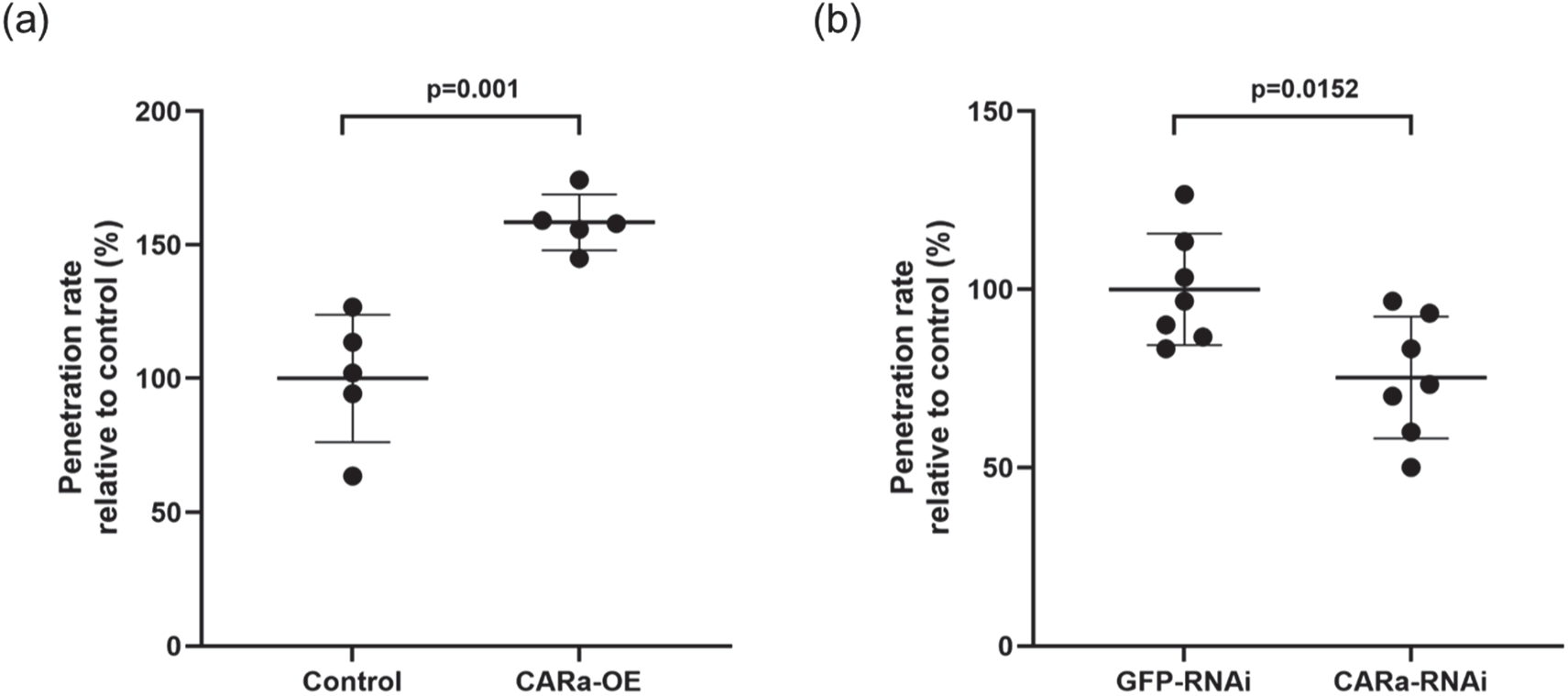
CARa affects *Blumeria hordei* penetration efficiency. The susceptibility of barley epidermal cells toward penetration by *Bh* was measured after transient overexpression (a) of CARa compared to pGY1 empty vector (control) or after knockdown (b) (CARa-RNAi) of *CARa* via RNAi compared to pIPKTA30N-GFP (GFP-RNAi). Each data point shows the penetration efficiency of a minimum of 50 plant–fungus interactions of a single experiment relative to its averaged control. RNAi-silencing specificity was confirmed using the si-Fi RNAi-off-target prediction tool (Luck et al., 2019, Figure S3). The mean values of all experiments are indicated with bars. Crossbars indicate the average susceptibility from minimum five independent biological experiments. Statistical differences were calculated with Tukey test, based on an ANOVA.

### CARa localizes at the extrahaustorial membrane

Host membrane reorganization takes place at microbial entry sites in plants (Dörmann et al., 2014). To investigate whether CARa shows specific or altered localization during fungal invasion, we transiently expressed GFP-tagged CARa (GFP-CARa) in barley epidermal cells and inoculated them with *Bh*. We co-expressed soluble mCherry as a cytoplasmic reference. At 20 hours post inoculation, we performed confocal laser scanning microscopy to assess subcellular protein localization. In non-attacked cell, the GFP-CARa signal was detectable in the cytoplasm and nucleoplasm similar to soluble mCherry (Figure 5a). By stark contrast, in penetrated cells, we recorded a GFP-CARa signal clearly distinct from that of cytosolic mCherry surrounding the haustorium (Figure 5b, for more examples see supplementary Figure S5). GFP-CARa exhibited a continuous, well-defined localization pattern surrounding the haustorium, while mCherry showed diffuse and variable fluorescence intensity. Notably, the CARa fluorescence showed uniform thickness across all observed confocal optical sections, indicating a specific EHM localization. To confirm that the observed GFP signal originated from GFP-CARa at the EHM and not from autofluorescence, we performed a λ-scan to record the spectrum of the emitted light. This spectral analysis confirmed that the signal at the haustorial neck and at the EHM derived specifically from GFP-CARa expression (emission peak around 510-520 nm; Figure 5 c,d). Cytosolic GFP-CARa signals were still detectable but weak when compared to the signal at the EHM. These findings suggest that CARa specifically localizes to the EHM during fungal invasion.

**Figure 5:**
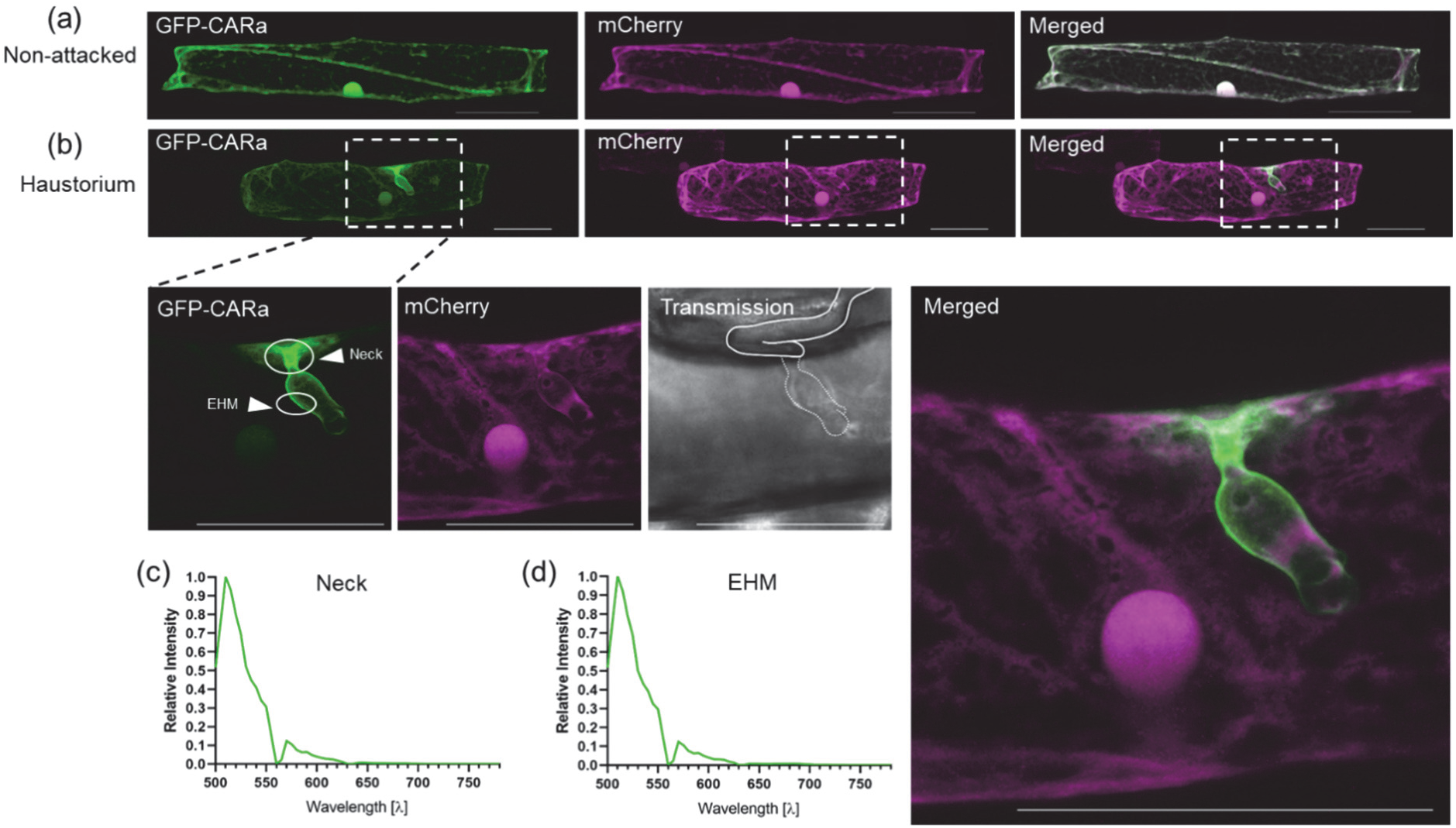
CARa localizes at the extrahaustorial membrane. (a) The subcellular localization of GFP-tagged CARa was investigated in barley via confocal laser scanning microscopy after transient transformation of epidermal cells. Free mCherry was co-transformed as a cytosolic marker. GFP-CARa showed a cytosolic localization. Images show Z-stack maximum intensity projections composed of at least 20 xy-optical sections captured in 1.5 µm Z-steps. Scale bar: 50 µm. Representative images from at least four independent biological experiments are shown. Image brightness was uniformly enhanced post-scanning for better visibility. (b) Images of an attacked haustorium containing cell were taken between 16 and 20 hours post inoculation. High-magnification images of haustorium are shown and arrows indicate the neck and the extrahaustorial membrane (EHM). (c) The circles in the GFP-CARa image show regions-of interest that were λ-scanned to evaluate fluorescence emission spectra. λ-scanning was performed in 5 nm detection steps after excitation with a 488 nm (GFP). Both, at EHM and neck emission spectra matched the typical GFP pattern.

## Discussion

To identify potential interactors of RIPb in barley, we performed a yeast-two hybrid (Y2H) screening using RIPb as a bait. Among the interacting partners obtained, we identified a C2-Domain Abscisic Acid-Related protein (CARa). The C2 domain, comprising 115-130 amino acid residues, was first identified in mammalian proteins kinase C (Nishizuka, 1988). In plants, C2 domains are present in various proteins, including synaptotagmins and phospholipases, and may appear as single or multiple domains, often in combination with other distinct domains within the same protein (Cui et al., 2023; Kopka et al., 1998). In Arabidopsis, CAR proteins possess a only single C2 domain and were suggested to interact with and recruit Pyrabactin resistance1 (PYR1)/PYR1-like (PYL)/Regulatory components of ABA receptors (RCAR) ABA receptors to the PM in a Ca^2+^-dependent manner and to promote membrane curvature (Diaz et al., 2016; Rodriguez et al., 2014). Structural and biochemical studies suggest that CAR proteins act as peripheral membrane components that protrude into the phospholipid bilayer and cluster at specific PM sites, serving as assembly hubs for ABA receptors (Diaz et al., 2016). Recently, studies have shown that the receptor-like kinase (RLK) FERONIA interacts with and phosphorylates CAR to modulate the nano-organization of the PM during FERONIA activation by the Rapid Alkalinization Factor 1 (RALF1) (Chen et al., 2023). This RALF1-FERONIA-CAR module plays an important role in regulating plant immunity at membrane nanodomains. CAR-bound Ca^2+^ ions interact with negatively charged phospholipids at the PM to induce membrane curvature (Chen et al., 2023; Diaz et al., 2016; Khan et al., 2019; Rodriguez et al., 2014).

Beyond their involvement in plant immunity, CAR proteins also contribute to abiotic stress responses. Qin et al. (2019) found that Lower Temperature 1 (LOT1) interacts with CAR9 and regulate its relocalization from the nucleus to the PM, where it interacts with ABA-receptors to regulate plant drought tolerance. Furthermore, CAR proteins were described as regulators of alkali stress tolerance via their interaction with the PM H^+^-ATPase AHA1 (Guo et al., 2024). A function was suggested earlier for ABA, RALFs, FERONIA and apoplastic pH in modulation of susceptibility to powdery mildew (Felle et al., 2004; Wiese et al., 2004; Kessler et al., 2010; Leicher et al., 2025). Crystallographic analysis of Arabidopsis CAR4 showed that it consists, similarly to the C2 domain of protein kinase C, of two four-stranded β-sheets that can coordinate calcium ions by aspartate-rich loops to bind phospholipids (Rodriguez et al., 2014). CAR4 possess aspartate residues in loop 1 (D34 and D39) and loop 3 (D85 and D87), that are conserved in barley CARa and responsible for coordinating Ca^2+^ to bridge C2 domain to phospholipids (Verdaguer et al., 1999). Mutation of these D85 and D87 in CAR4 (CAR4^D85A^ ^D87A^) and CAR1^D22A^ ^D27A^ abolished phosphatidylserine biding (Rodriguez et al., 2014). Furthermore, a rice CAR protein (OsGAP1 or OsCARe according to our phylogenetic analysis) binds the unconventional G-protein (OsYchF1) and serves as a GTPase-activating protein 1. OsCARe/OsGAP1 was reported as CAR4 homologue and possess a C2 domain (Yung et al., 2015). Crystal structure and site-directed mutagenesis show that OsGAP1 also contains conserved aspartate residues responsible for Ca^2+^ binding and phospholipid interaction. Phospholipid binding assays showed that mutation of D23A and D28A residues of OsGAP1 abolished the binding to phosphatidylserine, phosphatidylglycerol (PG) and cardiolipin (Yung et al., 2015). Additionally, Arabidopsis CAR6 (Enhanced Bending1, EHB1) was found to bind to PtdIns and PtdIns4P in the presence of Ca^2+^ (Khan et al., 2019). EHB1/CAR6 interacts with Iron-Regulated Transporter1 (IRT1) in a calcium-dependent way to inhibit its activity as a negative regulator of iron uptake, which contributes to iron homeostasis in plants. These findings support that CAR proteins are key regulators in multiple signalling pathways and plant stress responses (Cui et al., 2023). However, the exact mechanisms regulating these processes involving CAR remain unclear.

Barley CARa and other barley CAR proteins show conserved molecular signatures of typical CAR proteins and lipid binding (Figs. 1 and 2). PM and EHM lipid composition was recently associated with functions in susceptibility to accommodation of haustoria in intact plant epidermal cells cells (Qin et al., 2019; Weiß et al., 2025). We identified CARa as an interactor of RIPb, which itself interacts with RACB but not with the presumably inactive DNRACB. Overexpression of RIPb or its RACB-interacting coiled coil domain (RIPbCC2) promotes *Bh* invasion and haustoria accommodation in barley. RACB and RIPb both strongly accumulate at the haustorial neck region of developing haustoria, whereas their localisation at the EHM is weaker, though detectable for RIPb (McCollum et al. 2020). Barley RIPb has a potentially phospholipid-binding basic lysine-rich motif (KKGPK) at the carboxy-treminus, similar to AtRIP1/ICR1, where this motif mediates polar membrane association (S. Li et al., 2008). RACB also possesses a polybasic, anionic phospholipid-binding domain critical for its role in fungal invasion. Certain anionic phospholipids, including phosphoinositol-4-phosphate and phosphatidylserine, accumulate at *Bh* penetration sites (Weiß et al., 2025). These observations suggest that PM lipid composition changes during fungal attack, potentially recruiting membrane-associated proteins that function in plant-pathogen interaction. RACB, RIPb and CARa may be recruited to such sites or contribute to membrane domain formation by protein-protein-lipid interaction. CARa shares localization at the haustorial neck with RACB and RIPb but exhibits stronger association with the EHM, which is not observed for RACB or RIPb. The haustorial neck has been proposed to function in sorting PM-associated proteins for selective access to the EHM (Koh et al., 2005). It seems possible that RACB, RIPb, and CARa come together in lipid-specified PM domains at the site of pathogen attack, but the sorting into the EHM is more specific and preferably allows CARa to enter. CAR proteins itself may function in membrane curvature, ruffling or PM domain formation (Chen et al., 2023; Diaz et al., 2016). While we cannot confirm such function for barley CARa, its similarity to other CAR proteins with such functions makes it tempting to speculate that CARa could support membrane reorganizing for facilitated fungal invasion and EHM formation or influence EHM membrane identity. RACB was recently shown to interact with a barley phosphoinositide phospholipase C1 that also contains a C2-domain for membrane lipid targeting. This PLC1 protein acts in resistance (Weiß et al. 2025), potentially by competing for or removing lipids that otherwise would support fungal entry. Together, convergent evidence strongly supports the involvement of RACB, RACB-binding and phospholipid-binding proteins in modulating *Bh* entry into barley epidermal cells.

## Materials and methods

### Biological material and growth conditions

Wild-type barley (*Hordeum vulgare* L. cv. Golden Promise) was cultivated in a controlled growth chamber (Conviron, Winnipeg, Canada) at 18 °C with 65% relative humidity under a 16 h light/8 h dark photoperiod (150 µmol m⁻² s⁻¹). The powdery mildew fungus *Blumeria hordei* isolate A6 was propagated on Golden Promise plants under identical conditions and used for inoculations. Inoculation was performed by blowing conidiospores onto healthy plants or detached leaf segments enclosed in a plastic tent. Plants at 7-9 days post-inoculation were used as inoculum.

*Nicotiana benthamiana* was grown on a soil mixture containing five parts substrate and one part fine vermiculite (1-3 mm; Raiffeisen Gartenbau, Köln, Germany). Seeds were stratified for at least 2 days at 4 °C prior to transfer to long-day conditions (16 h light at 150 µmol m⁻² s⁻¹, 23 °C/8 h dark, 21 °C; 55% relative humidity).

### Cloning procedures

Barley *HvCARa* (HORVU.MOREX.r3.5HG0469960.1) cloning was performed using a combination of Gateway and classical cloning techniques. All oligonucleotides used bellow are listed in the table S4.

For Gateway cloning, the coding sequence of *HvCARa* was amplified by PCR from *Hordeum vulgare* cDNA using attB-flanked primers attB1-CARa_F and attB2-CARa_R. The amplicon was cloned into the pDONR223 vector using the BP Clonase^TM^ II (Invitrogen) reaction to generate the Entry clone. The *Hv*RIPb and *Hv*RIPbCC2 Entry clones were constructed using the same procedure, except that the templates were the plasmids pGEM-T easy-RIPb and pGEM-T easy-RIPbCC2 from McCollum et al. (2020), respectively,instead of cDNA. The primers used to amplify the insert were attB1-RIPb_F + attB1-RIPb_R, and attB1-RIPbCC2_F + attB1-RIPb_R respectively. In a next step, the inserts were transferred into the destination vectors through LR Clonase^TM^ II (Invitrogen) reaction to generate the final expression constructs.

For the FRET-FLIM, pDONR223-RIPb and pDONR223-RIPbCC2 were subcloned into the Gateway-compatible pGY1-meGFP destination vector (Engelhardt et al., 2023). pDONR223-CARa was recombined into Gateway-compatible pGY1-mCherry (Engelhardt et al., 2023). For the control constructs, pDONR223-GST-mCherry (Weiß et al., 2025) was cloned into Gateway pGY1 destination vector (Engelhardt et al., 2023) via LR reaction, and the pGY1-mCherry-MAGAP1 was generated by Hoefle et al., (2011).

For BiFC, nYFP-RIPb, was constructed in (McCollum et al., 2020). The other constructs generated in this study followed the same clonig strategy. nYFP-CARa and cYFP-CARa were amplified using the primers SpeI-CARa_F and SalI- CARa_R, and nYFPCC2 was amplified with SpeI+ RIPbCC2_F and SalI- RIPbCC2_R. The PCR products were digested with SpeI and SalI and ligated into pUC-SPYNE(R)173 and pUC-SPYCE(MR) plasmids (Waadt et al., 2008). The control constructs cYFP-DN RACB and cYFP-CA RACB were described by Schultheiss et al. (2008).

Cloning of pGADT7-CARa for the yeast two-hybrid (Y2H) assay was carried out via LR reaction using pDONR223-CARa as the Entry clone and pGADT7 (Clontech) as thedestination vector. pGADT7-RIPb, pGBKT7-RIPb, and pGBKT7-RACB WT were generated by McCollum et al. (2020).

For transient barley transformation, CARa was cloned using the same procedure described above and transferred via LR reaction into pGY1 destination vector (Engelhardt et al., 2023) to create the barley overexpression construct, and into pGY1-meGFP (Engelhardt et al., 2023) to generate the localization construct. RNAi-silencing construct for CARa was generated by first amplifying part of its coding sequences with the primers NotI-CARa-RNAi_F + SalI-CARa-RNAi_R. Fitting nucleotide sequence stretches with a high probability for gene-specific RNAi were predicted using the si-Fi software (Lück et al., 2019). PCR amplicon was inserted into pIPKTA38 vectors (Douchkov et al., 2005) in 3’-to-5’ orientation using SalI/NotI-mediated classical cloning (Thermo Fisher Scientific, Waltham, USA). From pIPKTA38, the sequence was transferred into the destination vector pIPKTA30N (Douchkov et al., 2005) via Gateway LR-reactions, creating antisense-hairpin-sense RNAi-construct.

For recombinant protein expression in *E. coli*, GST-CARa was amplified via PCR from pDONR223-CARa using SalI+CARa_F+ NotI+CARa_R primers, and ligated into pGEX-6P-1 (GE-Healthcare, Chicago, USA). via SalI/NotI-mediated classical cloning.

### Transient transformation of barley epidermal cells

Epidermal cells of 7-day-old primary leaves of wild-type barley cv. Golden Promise were transiently transformed by particle bombardment using a PDS-1000/HE system (Bio-Rad), following the protocol of (Schweizer et al., 1999). For each shot, 11 µl of a suspension of spherical gold particles suspension (27.5 µg/ml; 1 µm diameter; Bio-Rad, Hercules, USA) were coated with 1 µg of plasmid DNA encoding the constructs of interest and 0.5 µg of transformation markers (e.g. free fluorophores or GUS+). DNA-coated particles were prepared by adding CaCl₂ to a final concentration of 0.5 M and 3 µl of 2 mg/ml protamine (Sigma), followed by incubation for 30 min at room temperature. The particles were then washed sequentially with 70% (v/v) ethanol and 100% (v/v) ethanol, resuspended in 6 µl of absolute ethanol per shot, and loaded onto macrocarriers for bombardment.

### Penetration efficiency assay

Epidermal cells of transiently transformed barley leaves, either with plasmids for gene overexpression (pGY1; Schweizer et al., 1999) or RNAi-mediated silencing (pIPKTA30N-GFP; Douchkov et al., 2005), were inoculated with *Bh* 24 h after bombardment (overexpression) or 48 h after bombardment (RNAi). Penetration efficiency was assessed using a transient assay system based on a cytosolic GUS reporter (pUbiGUSPlus; Vickers et al., 2003; Addgene plasmid #64402, RRID:Addgene_64402), as previously described (Schweizer et al., 1999). We evaluated the penetration efficiency in five independent experiments, considering a minimum of 50 transformed GUS-expressing cells single cells attacked by the fungus per experiment.

For GUS staining, leaves were vacuum-infiltrated with staining solution (0.1 M Na₂HPO₄/NaH₂PO₄, pH 7.0; 0.01 M EDTA; 5 mM K₄[Fe(CN)₆]; 5 mM K₃[Fe(CN)₆]; 0.1% [v/v] Triton X-100; 20% [v/v] methanol; 0.5 mg ml⁻¹ X-Gluc), incubated for 4 h at 37 °C, and subsequently de-stained in 70% (v/v) ethanol for at least 24 h. Transformed cells were identified by light microscopy, and fungal structures were stained with Calcofluor White solution (0.3% [w/v] Calcofluor White M2R, F-3543, Sigma-Aldrich, St. Louis, USA; 50 mM Tris; 2% [v/v] Tween-20; pH 9). A successful penetration event was scored when a haustorium was present. For each construct, at least 50 plant–fungus interactions were analyzed per replicate, using empty vector–transformed cells as controls, and a minimum of five independent biological replicates was performed.

RNAi construct efficiency was validated prior to susceptibility assays by transient co-expression of GFP-CARa with cytosolic mCherry as a transformation marker. GFP-CARa was cotransformed either with the empty vector pIPKTA30N (control) or with a CARa-RNAi hairpin construct targeting amino acids 132-361. Confocal z-stack images were taken 48 h after bombardment, and fluorescence intensities of GFP and mCherry were quantified. RNAi efficiency was calculated as the ratio of mean mCherry-normalized GFP fluorescence in CARa-silenced cells relative to mCherry-normalized GFP-CARa control cells.

### Confocal laser scanning microscopy

For subcellular localization experiments, transiently transformed barley leaves expressing fluorophore fusion proteins were inoculated with *Bh* 6-8 h after bombardment. Images were acquired 16-40 h after inoculation using a Leica TCS SP5 confocal laser scanning microscope. CFP was excited with a 458 nm Argon laser line and detected at 463-485 nm; GFP was excited with a 488 nm Argon laser line and detected at 500–550 nm; YFP with a 514 nm Argon laser line and detected at 525–570 nm; and mCherry with a 561 nm DPSS diode laser and detected at 570-620 nm. Highly fluorescent samples were recorded using photomultipliers (PMTs), whereas samples with low fluorescence were analyzed with hybrid detectors (HyDs;Leica Microsystems). Barley epidermal cells were imaged by sequential scanning as z-stacks of single XY optical sections, with z-step sizes specified in the figure legends.

For λ-scanning, GFP was excited with a 488 nm Argon laser line, and emission spectra were collected in xyλ-mode at 5 nm intervals across 500–780 nm. Spectra were extracted from regions of interest showing strong fluorescence and analyzed using Leica LAS X software (v3.5.1). Normalized mean fluorescence intensities were exported and plotted in GraphPad Prism v8.0 (GraphPad Software, San Diego, USA).

### Bimolecular fluorescence complementation (BiFC)

BiFC assays were performed in barley epidermal cells transiently transformed with constructs encoding N- and C-terminal fragments of YFP (pUC-SPYNE and pUC-SPYCE) fused to proteins of interest. Co-expression of a cytosolic mCherry construct was included as a transformation and normalization control. Leaves were imaged 24 h after particle bombardment using confocal laser scanning microscopy under identical acquisition settings (see above). Fluorescence was quantified in Fiji (ImageJ; Schindelin et al., 2012) by measuring the mean fluorescence intensity (MFI) across the entire transformed cell in both YFP and mCherry channels. The BiFC signal was expressed as the ratio of YFP to mCherry MFI (MFI_YFP/MFI_mCherry) to correct for variation in transformation efficiency and expression levels. For each construct, a minimum of 25 cells were analyzed per independent experiment, and at least two biological replicates were performed.

### FRET-FLIM measurements

FRET-FLIM experiments were performed in transiently transformed barley epidermal cells using an FCS/FLIM-FRET/rapidFLIM upgrade kit (PicoQuant, Berlin, Germany) mounted on an Olympus FV3000 confocal microscope on an IX63 stand (Olympus, Tokyo, Japan). Cells were imaged with a UPLSAPO 60XW 60×/NA 1.2 water immersion objective with 4× zoom to meet the Nyquist criterion. GFP was excited with a 485 nm pulsed diode laser (pulse rate 40 MHz; LDH-D-C-485), and fluorescence emission was detected using Hybrid PMA 40 detectors coupled to TimeHarp 260 TCSPC modules (PicoQuant) with 25 ps resolution. For each measurement, ≥500 photons were collected in the brightest pixel. Regions of interest (ROIs) were selected from focused areas showing visible fluorescence. Lifetime decay fitting was performed in SymPhoTime 64 software (v2.6; PicoQuant) using n-exponential reconvolution (n = 2) with instrument response correction. Only measurements with χ² values between 1.0 and 2.0 and positive amplitudes were considered. GFP and mCherry emission were collected between 500–540 nm and 580–620 nm, respectively. At least 10–15 cells per interaction were analyzed in three independent replicates. Intensity-weighted average GFP lifetimes (τ) were plotted in GraphPad Prism v8.0 (GraphPad Software, San Diego, USA).

### Yeast Two-Hybrid Assays

Yeast strain Y2HGold was co-transformed with bait (pGBKT7) and prey (pGADT7) constructs following the Yeastmaker™ Yeast Transformation System 2 protocol (Clontech). Transformed colonies were spotted on complete supplement medium lacking leucine and tryptophan (CSM–LW) to confirm successful transformation. For interaction analysis, colonies were spotted on selective medium lacking leucine, tryptophan, adenine, and histidine (CSM–LWAH) supplemented with 2.5 mM 3-amino-1,2,4-triazole (3-AT) and incubated at 30 °C for at least 3 days. Growth on CSM–LW confirmed transformation, while growth on CSM–LWAH + 3-AT indicated reporter gene activation and putative protein–protein interactions. A positive control for interaction was murine p53 with SV40 large T-antigen (B. Li & Fields, 1993). Protein extraction from yeast cells were prepared for Western blot verification of bait and prey expression.

### RNA Extraction and RT-qPCR

For gene expression barley cv. Golden Promise plants were sampled in four biological replicates. Epidermal peels from inoculated (24 h after *Bh* infection) or uninoculated leaves were excised and immediately frozen in liquid nitrogen. Total RNA was extracted using TRIzol reagent (Invitrogen) and purified with the Direct-zol^TM^ RNA purification Kit (Zymo Research) following the manufacturer’s instructions, followed by DNase I digestion. For cDNA synthesis, 2 µg of RNA per sample were reverse transcribed using the RevertAid RT Kit (Thermo Fisher Scientific, Waltham, USA), and the resulting cDNA was diluted 1:5 prior to analysis.

RT-qPCR was performed on 3 µl of diluted cDNA using the Maxima SYBR Green/ROX qPCR Master Mix (2×; Thermo Fisher Scientific) in 20 µl reactions with gene-specific primers (Table S4). Amplification was carried out on an AriaMx3000 system (Agilent) under the following conditions: 40 cycles of 95 °C for 10 s and 60 °C for 15 s, followed by a melting curve analysis from 65–95 °C. *HvUbiquitin2* (HvUBC2_fwd TCTCGTCCCTGAGATTGCCCACAT; HvUBC2_rev TTTCTCGGGACAGCAACACAATCTTCT) served as the internal reference gene (Schnepf et al., 2018). HvCARa was amplified using primers: qPCR_CARa_2_fwd: GCCATCAGGGACTTCACCTC andqPCR_CARa_2_rev:AAGGTTCCTTGACGGAGAAGAC. Relative expression levels were calculated using the 2^−ΔΔCt^ method (Livak & Schmittgen, 2001).

### Production and purification of recombinant proteins

For lipid-binding assays, recombinant proteins were expressed in *E. coli* Rosetta 2 cells transformed with pGEX-6P-1 based plasmid carrying the construct of interest (see Molecular Cloning). Single colonies were used to inoculate 30 ml 2YT starter cultures (10 g/L yeast extract, 16 g/L tryptone, 5 g/L NaCl, 2 g/L D-glucose) and grown overnight at 18 °C with shaking. The following day, 300 ml of fresh 2YT were inoculated with 1% of the starter culture and grown at 37 °C until OD_600_ reached 0.6–0.8. Protein expression was induced with 0.1 mM IPTG after a 30 min cooling step on ice, and cultures were incubated for 18–20 h at 18 °C with shaking.

Cells were harvested by centrifugation at 3000 g, aliquoted in 50 ml fractions, and frozen in liquid nitrogen, and stored at –80 °C. Cell pellets were resuspended in 3 ml extraction buffer and incubated on ice for 30–45 min. Proteins were extracted in 50 mM Tris–HCl pH 7.4, 150 mM NaCl, 1 mM DTT, 2 mg/ml lysozyme, and SigmaFast protease inhibitor (EDTA-free; Sigma-Aldrich). Cells were lysed byheat shock,repeatedly freezing in liquid nitrogen for 1 minute and thawing threetimes. Supernatants were used for affinity purification.

GST fusions were enriched on Pierce™ glutathione agarose (Thermo Fisher Scientific). Resin was packed into Pierce™ centrifugation columns (ThermoFisher) and washed with equilibration buffer (GST: 50 mM Tris–HCl pH 7.4, 150 mM NaCl, 1 mM DTT). Crude extracts were incubated with resin for 1 h at room temperature with gentle tumbling, followed by three washes with equilibration buffer. Fusion proteins were eluted in GST buffer supplemented with 50 mM gluthatione reduced (Duchefa Biochemie). Eluates were stored at –80 °C until further use.

### Protein–lipid-overlay experiments

Protein–lipid interactions were assessed using commercially available PIP Strips (Echelon Biosciences Inc., MoBiTec GmbH, Gottingen, Germany). Membranes were blocked for 1 hour in PBS containing Tween-20 (10 mM phosphate; 137 mM NaCl; 2.7 mM KCl) and 3% (w/v) bovine serum albumin (BSA) fatty acid free powder (Sigma). Membranes were then incubated overnight at 4 °C with 5 µg/ml purified protein in the same blocking solution supplemented with 10µM CaCl2, under gentle agitation in the dark. Following incubation, membranes were washed three times with PBS and probed with anti-GST antibody (Chromotek) diluted in 3% BSA/PBS for 2 hours at room temperature. After three additional washes, membranes were incubated with the secondary antibody anti-rat-AP (Sigma) for one hour under the same conditions. Washing was completed with two rinses in PBS and one in AP buffer (100 mM Tris–HCl, pH 9.5; 100 mM NaCl; 5 mM MgCl₂).

Protein binding was visualized by adding 0.175 mg/ml BCIP (5-bromo-4-chloro-3-indolyl phosphate disodium salt; Roth, Karlsruhe, Germany) and 0.338 mg/ml NBT (nitro blue tetrazolium chloride; Roth) in AP buffer. The reaction was stopped by rinsing with distilled water once sufficient color development was observed.

### Alignments and Phylogenetic Analysis

Protein Sequences of *Hv*CAR (*Hordeum vulgare*), *At*CAR (*Arabidopsis thaliana*), *Os*CAR (*Oryza sativa* ssp. *japonica*), and *Bd*CAR (*Brachypodium distachyon*) were retrieved from EnsemblPlants (https://plants.ensembl.org/index.html). Multiple sequence alignment was performed with ClustalO (https://www.ebi.ac.uk/Tools/msa/clustalo/) and visualized in Jalview (v2.10.5). Phylogenetic analysis was conducted using MAFFT (Katoh & Standley, 2013), and the phylogenetic tree was inferred using IQ-TREE (Nguyen et al., 2015) using the best-fit substitution model automatically selected. The resulting tree was visualized using iTOL (https://itol.embl.de/; (Letunic & Bork, 2021). Protein sequences used in the analysis are listed in supplementary file table S1 and S2.

### Statistical analysis

Unpaired two-sided Student’s *t*-tests were used for pairwise comparisons. For multiple group comparisons, one-way ANOVA followed by Tukey’s post-hoc test was applied. All statistical analyses were carried out using GraphPad Prism (GraphPad Software, San Diego, USA). Differences were considered significant at *P* < 0.05.

## Supporting information

are available online

## Acknowledgements

We are grateful to Marion Müller (TU Munich) for support of the phylogenetic analyses and to Veronika Schmitt (TU Munich) for technical support.

## Author contributions

MB and RH designed the study. CB and CM carried out experiments. RH acquired funding. MB prepared the figures. MB and RH wrote the manuscript. CM read and approved the manuscript.

Supplementary figures and tables are available online.

