## Supplementary material for "Barley C2-Domain Abscisic Acid-Related protein CARa supports susceptibility to *Blumeria hordei* and localizes to the extrahaustorial membrane": are available online

### Supplementary Materials

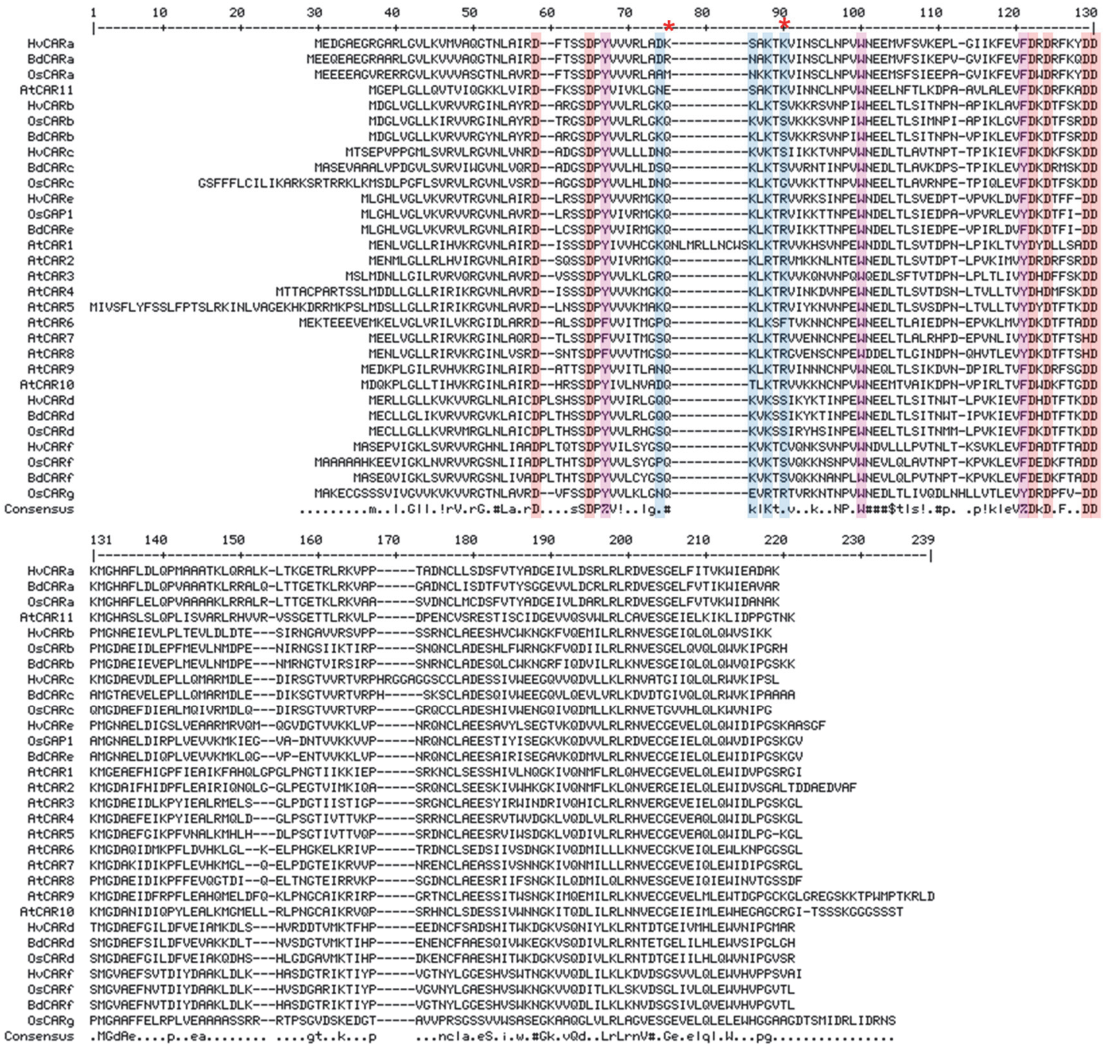

**Figure S1:** Alignment of *H. vulgare*, *A. thaliana*, *O. sativa* and *B. distachyon* as basis for phylogenetic tree in Figure 1. Conserved amino acid residues mentioned in text are highlighted.

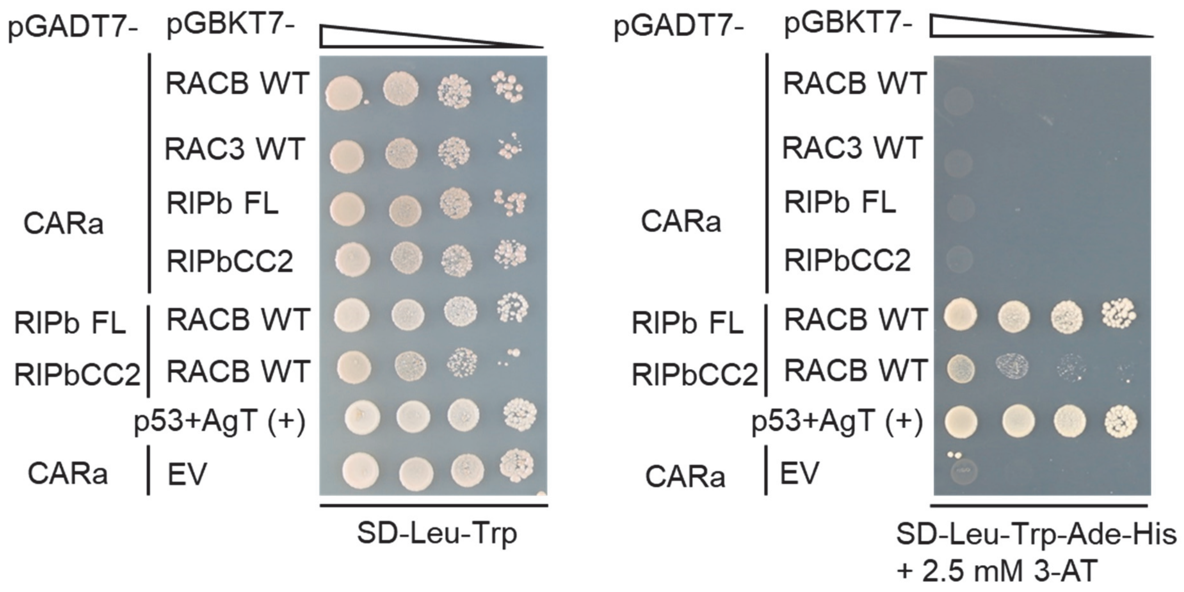

**Figure S2:** CARa protein did not interact with RIPb, and RIPbCC2 in yeast-two-hybrid. **Yeast two-hybrid assay** was performed using binding domain (BD)-tagged RACB wild type, RIPb or RIPbCC2 and activation domain (AD)-tagged CARa, RIPb or RIPbCC2 expressed in yeast. Decreasing numbers of cells (10 fold dilutions) were grown on transformation-selective plates lacking leucine and tryptophan, as well as on interaction-selective plates lacking leucine, tryptophan, histidine, and adenine supplemented with 2.5 mM 3-aminotriazole. Growth indicating protein interaction was observed for RIPb, and RIPbCC2 with RACB and between T-antigen and p53 that served as a positive control.

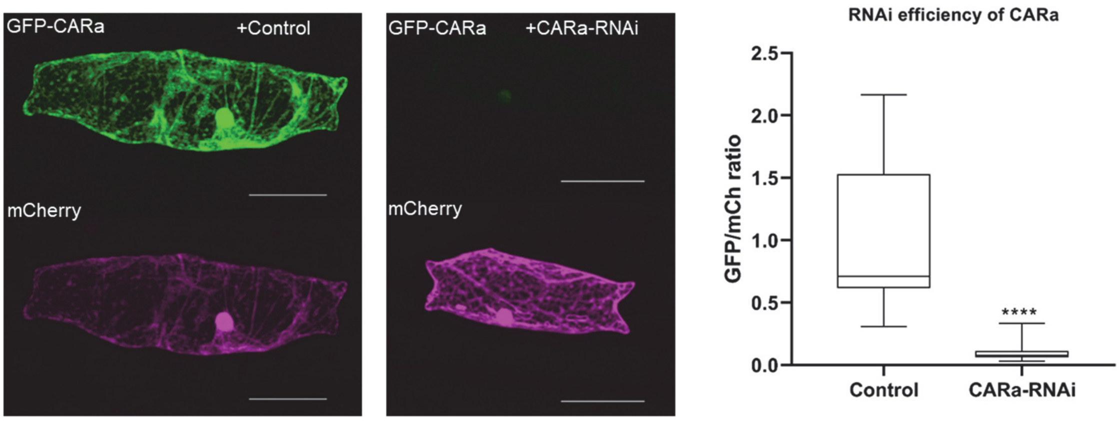

**Figure S3:** RNAi efficiency of CARa hairpin construct in transient transformation of barley epidermal cells. Epidermal cells from 7-day-old barley primary leaves were transiently transformed using particle bombardment with constructs for overexpressing a GFP-tagged CARa, either alone or in combination with an RNAi silencing construct. A cytosolic mCherry expression construct was co-delivered to monitor transformation efficiency and enable ratiometric fluorescence quantification. Microscopy images represent maximum intensity projections of at least 15 optical sections acquired at 2 µm intervals. Scale bar = 50 µm. Boxes displays the GFP fluorescence expressed as a ratio to mCherry fluorescence per transformed cell, with the entire cell area used as the region of interest for intensity measurements. Each dot corresponds to an individual cell. Statistical significance was assessed using a t-test ($p<0.0001$).

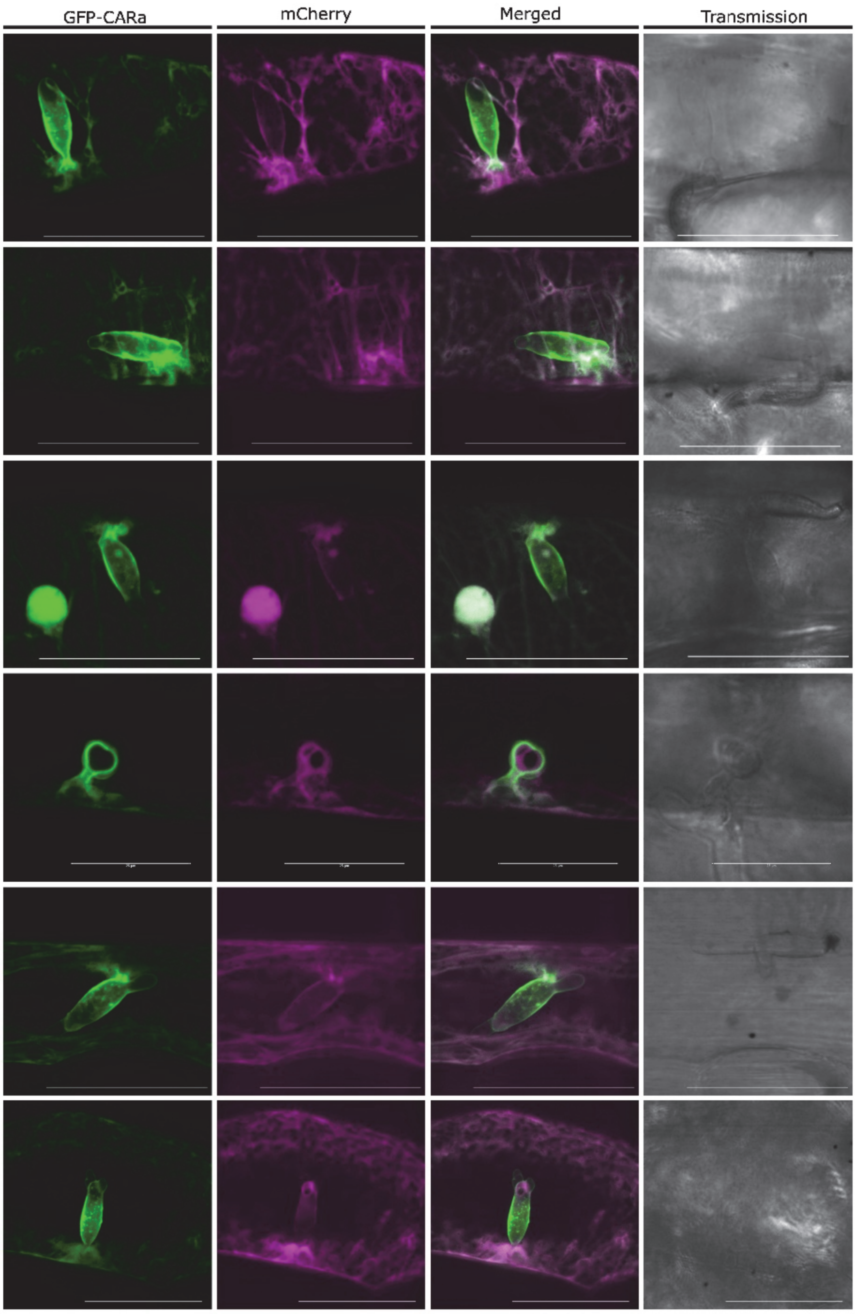

**Figure S4:** CARa localizes at the extrahaustorial membrane. (a) The subcellular localization of GFP-tagged CARa was investigated in barley via confocal laser scanning microscopy after transient transformation of epidermal cells. Free mCherry was co-transformed as a cytosolic marker. GFP-CARa showed a cytosolic localization. Images show Z-stack maximum intensity projections composed of at least 20 xy-optical sections captured in 1.5 µm Z-steps. Scale bar: 50 µm. Six examples of cells represent typical images from at least four independent experiments. Image brightness was uniformly enhanced post-scanning for better visibility.

**Supplementary Tables; Bradai et al.**

**Table S1:** List of genes encoding CARs in barley, rice and Brachypodium with gene identifiers, and protein sequence length

| *Hordeum vulgare* | | | *Oryza sativa* | | | *Brachypodium distachyon* | | |
|---|---|---|---|---|---|---|---|---|
| Gene name | Gene ID | Protein size (aa) | Gene name | Gene ID | Protein size (aa) | Gene name | Gene ID | Protein size (aa) |
| *Hv*CARa | HORVU.MOREX.r3.5 HG0469960.1 | 171 | *Os*CARa | Os09t0251800 | 171 | *Bd*CARa | BRADI_4g08660v3 | 172 |
| *Hv*CARb | HORVU.MOREX.r3.2 HG0156770.1 | 161 | *Os*CARb | Os07t0108400 | 163 | *Bd*CARb | BRADI_1g59226v3 | 164 |
| *Hv*CARc | HORVU.MOREX.r3.2 HG0156820.1 | 170 | *Os*CARc | Os07t0108500 | 183 | *Bd*CARc | BRADI_1g59232v3 | 171 |
| *Hv*CARd | HORVU.MOREX.r3.6 HG0630040.1 | 166 | *Os*CARd / OsSTA1 96 | Os07t0501700 | 166 | *Bd*CARd | BRADI_1g26840v3 | 166 |
| *Hv*CARe | HORVU.MOREX.r3.2 HG0202460.1 | 169 | *Os*CARe / OsGAP1 | Os02t0327000 | 165 | *Bd*CARe | BRADI_3g10810v3 | 165 |
| *Hv*CARf | HORVU.MOREX.r3.4 HG0418640.1 | 169 | *Os*CARf | Os07t0500300 | 173 | *Bd*CARf | BRADI_1g26850v3 | 168 |
| | | | *Os*CARg | Os07t0462500 | 185 | | | |

**Table S2:** Percent identity matrix of CAR protein from *Arabidopsis thaliana, Hordeum vulgare, Oryza sativa and Brachypodium distachyon.* Color code ranges from low (blue) to high (red) amino acid identiry of Clustal_Omega aligned proteins. Gene IDs: see table S1 and for Arabidopsis: AtCAR1, At5g37740; AtCAR2, At1g66360; AtCAR3, At1g73580; AtCAR4, At3g17980; AtCAR5, At1g48590; AtCAR6, At1g70800; AtCAR7, At1g70810; AtCAR8, At1g23140; AtCAR9, At1g70790; AtCAR10, At2g01540; AtCAR1, At5g47710

| | AtCAR11 | OsCARa | HvCARa | BdCARa | OsCARg | HvCARf | OsCARf | BdCARf | HvCARd | OsCARd | BdCARd | OsCARb | HvCARb | BdCARb | OsCARc | HvCARc | BdCARc | HvCARe | OsGAP1 | BdCARe | AtCAR3 | AtCAR4 | AtCAR5 | AtCAR1 | AtCAR2 | AtCAR8 | AtCAR6 | AtCAR7 | AtCAR9 | AtCAR10 |
|---|---|---|---|---|---|---|---|---|---|---|---|---|---|---|---|---|---|---|---|---|---|---|---|---|---|---|---|---|---|---|
| AtCAR11 | 100 | 51,83 | 52,44 | 52,44 | 35 | 35,37 | 36,59 | 35,37 | 34,76 | 36,59 | 37,2 | 36,81 | 39,75 | 40,24 | 39,75 | 40,74 | 40,24 | 42,68 | 46,01 | 44,17 | 42,68 | 40,85 | 39,02 | 39,16 | 40,61 | 37,2 | 36,59 | 39,02 | 40,96 | 37,95 |
| OsCARa | 51,83 | 100 | 77,78 | 79,53 | 35,15 | 32,32 | 33,14 | 33,54 | 34,57 | 33,33 | 33,33 | 38,89 | 40,37 | 38,89 | 33,93 | 35,15 | 35,5 | 39,88 | 40,74 | 40,12 | 37,58 | 38,46 | 39,64 | 37,2 | 37,42 | 33,95 | 34,91 | 35,19 | 38,18 | 38,79 |
| HvCARa | 52,44 | 77,78 | 100 | 83,63 | 34,55 | 34,15 | 34,91 | 34,76 | 35,19 | 33,95 | 33,95 | 38,27 | 39,75 | 37,65 | 35,71 | 36,36 | 35,5 | 40,49 | 39,51 | 40,74 | 38,18 | 38,46 | 39,64 | 37,2 | 38,65 | 33,95 | 34,32 | 35,8 | 40 | 41,82 |
| BdCARa | 52,44 | 79,53 | 83,63 | 100 | 33,94 | 32,32 | 32,54 | 32,32 | 35,19 | 33,95 | 34,57 | 39,51 | 39,13 | 37,65 | 34,91 | 35,76 | 34,32 | 40,49 | 40,12 | 41,36 | 35,76 | 35,88 | 37,65 | 35,98 | 36,2 | 35,19 | 32,35 | 34,57 | 41,21 | 40,61 |
| OsCARg | 35 | 35,15 | 34,55 | 33,94 | 100 | 39,75 | 40 | 40,62 | 39,24 | 37,97 | 42,41 | 40,76 | 41,94 | 41,14 | 39,51 | 40,99 | 43,03 | 45,45 | 45,96 | 43,48 | 43,21 | 46,39 | 44,85 | 43,48 | 43,29 | 38,36 | 39,16 | 40,88 | 42,11 | 42,17 |
| HvCARf | 35,37 | 32,32 | 34,15 | 32,32 | 39,75 | 100 | 77,98 | 84,52 | 47,59 | 50,6 | 50,6 | 46,63 | 45,96 | 46,95 | 46,01 | 45,73 | 42,77 | 45,12 | 42,94 | 39,88 | 39,52 | 42,51 | 41,57 | 43,64 | 40,85 | 42,07 | 38,55 | 40,24 | 41,57 | 36,75 |
| OsCARf | 36,59 | 33,14 | 34,91 | 32,54 | 40 | 77,98 | 100 | 87,5 | 53,61 | 56,63 | 55,42 | 49,08 | 49,07 | 48,17 | 46,43 | 49,09 | 42,11 | 44,17 | 42,59 | 41,98 | 41,92 | 40,35 | 42,11 | 43,29 | 44,17 | 39,26 | 41,18 | 41,72 | 42,42 | 41,82 |
| BdCARf | 35,37 | 33,54 | 34,76 | 32,32 | 40,62 | 84,52 | 87,5 | 100 | 51,81 | 54,82 | 53,61 | 49,08 | 49,07 | 50 | 47,85 | 49,39 | 42,77 | 44,17 | 43,21 | 41,98 | 42,77 | 42,17 | 43,37 | 42,68 | 44,17 | 41,72 | 41,82 | 42,33 | 43,03 | 41,21 |
| HvCARd | 34,76 | 34,57 | 35,19 | 35,19 | 39,24 | 47,59 | 53,61 | 51,81 | 100 | 79,52 | 78,31 | 51,53 | 50,93 | 50 | 49,69 | 49,38 | 40,24 | 48,77 | 49,07 | 49,69 | 42,68 | 45,73 | 47,56 | 43,29 | 47,85 | 41,72 | 42,94 | 45,4 | 42,68 | 39,63 |
| OsCARd | 36,59 | 33,33 | 33,95 | 33,95 | 37,97 | 50,6 | 56,63 | 54,82 | 79,52 | 100 | 78,31 | 54,6 | 54,66 | 53,66 | 50,93 | 50 | 42,07 | 53,09 | 52,17 | 50,93 | 45,12 | 48,17 | 50,61 | 45,73 | 47,24 | 46,01 | 44,17 | 47,85 | 44,51 | 41,46 |
| BdCARd | 37,2 | 33,33 | 33,95 | 34,57 | 42,41 | 50,6 | 55,42 | 53,61 | 78,31 | 78,31 | 100 | 56,44 | 53,42 | 55,49 | 50,93 | 53,7 | 45,12 | 54,32 | 54,66 | 54,66 | 46,34 | 48,78 | 51,22 | 45,12 | 48,47 | 42,94 | 44,17 | 47,24 | 44,51 | 41,46 |
| OsCARb | 36,81 | 38,89 | 38,27 | 39,51 | 40,76 | 46,63 | 49,08 | 49,08 | 51,53 | 54,6 | 56,44 | 100 | 76,4 | 81,6 | 59,01 | 56,79 | 48,47 | 55,28 | 55,62 | 55 | 54,6 | 52,76 | 53,37 | 50,92 | 47,53 | 53,09 | 50 | 53,7 | 53,37 | 51,53 |
| HvCARb | 39,75 | 40,37 | 39,75 | 39,13 | 41,94 | 45,96 | 49,07 | 49,07 | 50,93 | 54,66 | 53,42 | 76,4 | 100 | 82,61 | 57,76 | 55,9 | 52,17 | 56,6 | 57,59 | 56,96 | 52,17 | 54,04 | 51,55 | 50,31 | 47,5 | 53,12 | 48,12 | 50,63 | 52,8 | 48,45 |
| BdCARb | 40,24 | 38,89 | 37,65 | 37,65 | 41,14 | 46,95 | 48,17 | 50 | 50 | 53,66 | 55,49 | 81,6 | 82,61 | 100 | 59,63 | 58,02 | 53,66 | 58,02 | 59,01 | 57,76 | 54,88 | 53,05 | 49,39 | 50 | 47,85 | 52,76 | 47,24 | 49,69 | 54,27 | 50 |
| OsCARc | 39,75 | 33,93 | 35,71 | 34,91 | 39,51 | 46,01 | 46,43 | 47,85 | 49,69 | 50,93 | 50,93 | 59,01 | 57,76 | 59,63 | 100 | 73,78 | 63,69 | 53,75 | 54,09 | 57,23 | 51,22 | 49,13 | 48,63 | 47,2 | 47,5 | 50 | 44,38 | 52,5 | 51,85 | 46,91 |
| HvCARc | 40,74 | 35,15 | 36,36 | 35,76 | 40,99 | 45,73 | 49,09 | 49,39 | 49,38 | 50 | 53,7 | 56,79 | 55,9 | 58,02 | 73,78 | 100 | 73,94 | 49,69 | 50 | 51,25 | 48,48 | 47,88 | 47,27 | 41,36 | 46,58 | 39,75 | 40,24 | 44,1 | 49,08 | 44,79 |
| BdCARc | 40,24 | 35,5 | 35,5 | 34,32 | 43,03 | 42,77 | 42,11 | 42,77 | 40,24 | 42,07 | 45,12 | 48,47 | 52,17 | 53,66 | 63,69 | 73,94 | 100 | 48,47 | 47,53 | 45,68 | 44,31 | 43,86 | 42,69 | 42,07 | 41,1 | 39,88 | 37,65 | 42,33 | 47,27 | 39,39 |
| HvCARe | 42,68 | 39,88 | 40,49 | 40,49 | 45,45 | 45,12 | 44,17 | 44,17 | 48,77 | 53,09 | 54,32 | 55,28 | 56,6 | 58,02 | 53,75 | 49,69 | 48,47 | 100 | 78,18 | 79,39 | 51,83 | 59,15 | 59,51 | 56,97 | 52,38 | 52,15 | 48,78 | 56,44 | 49,7 | 46,15 |
| OsGAP1 | 46,01 | 40,74 | 39,51 | 40,12 | 45,96 | 42,94 | 42,59 | 43,21 | 49,07 | 52,17 | 54,66 | 55,62 | 57,59 | 59,01 | 54,09 | 50 | 47,53 | 78,18 | 100 | 86,67 | 50,31 | 58,9 | 59,88 | 53,66 | 51,83 | 52,47 | 50,92 | 58,02 | 51,52 | 49,7 |
| BdCARe | 44,17 | 40,12 | 40,74 | 41,36 | 43,48 | 39,88 | 41,98 | 41,98 | 49,69 | 50,93 | 54,66 | 55 | 56,96 | 57,76 | 57,23 | 51,25 | 45,68 | 79,39 | 86,67 | 100 | 51,53 | 57,06 | 58,02 | 56,71 | 53,66 | 51,85 | 52,76 | 59,26 | 56,36 | 52,12 |
| AtCAR3 | 42,68 | 37,58 | 38,18 | 35,76 | 43,21 | 39,52 | 41,92 | 42,77 | 42,68 | 45,12 | 46,34 | 54,6 | 52,17 | 54,88 | 51,22 | 48,48 | 44,31 | 51,83 | 50,31 | 51,53 | 100 | 69,05 | 62,87 | 58,79 | 55,49 | 50,61 | 49,1 | 57,32 | 57,83 | 51,2 |
| AtCAR4 | 40,85 | 38,46 | 38,46 | 35,88 | 46,39 | 42,51 | 40,35 | 42,17 | 45,73 | 48,17 | 48,78 | 52,76 | 54,04 | 53,05 | 49,13 | 47,88 | 43,86 | 59,15 | 58,9 | 57,06 | 69,05 | 100 | 78,98 | 64,85 | 59,15 | 53,66 | 47,4 | 57,93 | 56,63 | 51,2 |
| AtCAR5 | 39,02 | 39,64 | 39,64 | 37,65 | 44,85 | 41,57 | 42,11 | 43,37 | 47,56 | 50,61 | 51,22 | 53,37 | 51,55 | 49,39 | 48,63 | 47,27 | 42,69 | 59,51 | 59,88 | 58,02 | 62,87 | 78,98 | 100 | 61,59 | 57,67 | 56,44 | 50 | 58,9 | 55,15 | 52,73 |
| AtCAR1 | 39,16 | 37,2 | 37,2 | 35,98 | 43,48 | 43,64 | 43,29 | 42,68 | 43,29 | 45,73 | 45,12 | 50,92 | 50,31 | 50 | 47,2 | 41,36 | 42,07 | 56,97 | 53,66 | 56,71 | 58,79 | 64,85 | 61,59 | 100 | 67,66 | 53,94 | 55,15 | 60,61 | 55,09 | 56,29 |
| AtCAR2 | 40,61 | 37,42 | 38,65 | 36,2 | 43,29 | 40,85 | 44,17 | 44,17 | 47,85 | 47,24 | 48,47 | 47,53 | 47,5 | 47,85 | 47,5 | 46,58 | 41,1 | 52,38 | 51,83 | 53,66 | 55,49 | 59,15 | 57,67 | 67,66 | 100 | 49,39 | 51,22 | 55,49 | 50,87 | 49,13 |
| AtCAR8 | 37,2 | 33,95 | 33,95 | 35,19 | 38,36 | 42,07 | 39,26 | 41,72 | 41,72 | 46,01 | 42,94 | 53,09 | 53,12 | 52,76 | 50 | 39,75 | 39,88 | 52,15 | 52,47 | 51,85 | 50,61 | 53,66 | 56,44 | 53,94 | 49,39 | 100 | 60 | 69,09 | 53,33 | 49,7 |
| AtCAR6 | 36,59 | 34,91 | 34,32 | 32,35 | 39,16 | 38,55 | 41,18 | 41,82 | 42,94 | 44,17 | 44,17 | 50 | 48,12 | 47,24 | 44,38 | 40,24 | 37,65 | 48,78 | 50,92 | 52,76 | 49,1 | 47,4 | 50 | 55,15 | 51,22 | 60 | 100 | 74,55 | 57,23 | 55,42 |
| AtCAR7 | 39,02 | 35,19 | 35,8 | 34,57 | 40,88 | 40,24 | 41,72 | 42,33 | 45,4 | 47,85 | 47,24 | 53,7 | 50,63 | 49,69 | 52,5 | 44,1 | 42,33 | 56,44 | 58,02 | 59,26 | 57,32 | 57,93 | 58,9 | 60,61 | 55,49 | 69,09 | 74,55 | 100 | 61,21 | 60 |
| AtCAR9 | 40,96 | 38,18 | 40 | 41,21 | 42,11 | 41,57 | 42,42 | 43,03 | 42,68 | 44,51 | 44,51 | 53,37 | 52,8 | 54,27 | 51,85 | 49,08 | 47,27 | 49,7 | 51,52 | 56,36 | 57,83 | 56,63 | 55,15 | 55,09 | 50,87 | 53,33 | 57,23 | 61,21 | 100 | 61,67 |
| AtCAR10 | 37,95 | 38,79 | 41,82 | 40,61 | 42,17 | 36,75 | 41,82 | 41,21 | 39,63 | 41,46 | 41,46 | 51,53 | 48,45 | 50 | 46,91 | 44,79 | 39,39 | 46,15 | 49,7 | 52,12 | 51,2 | 51,2 | 52,73 | 56,29 | 49,13 | 49,7 | 55,42 | 60 | 61,67 | 100 |

OsGAP1= OsCARe

**Table S3:** Tissue-specific *Hv*CAR gene expression based on RNAseq (James Hutton Institute) in fragments per reads per kilobase million (FPKM). Intensity of green colour indicates higher values, light green shows low values, and white fields for no expression.

| | FPKM | | | | | | | | | | | | | | | |
|---|---|---|---|---|---|---|---|---|---|---|---|---|---|---|---|---|
| **Input** | 4-day embryos dissected from germinating grains | Etiolated seedling (10 days) | Shoots from the seedlings (10 cm shoot stage) | Peeled epidermis (4 weeks) | Roots from the seedlings (10 cm shoot stage) | Root (4 weeks) | Developing tillers at six leaf stage, 3rd internode | Rachis (5 weeks pa) | Young developing inflorescences (5mm) | Developing inflorescences (1-1.5 cm) | Developing grain, bracts removed (5 DPA) | Developing grain, bracts removed (15 DPA) | Lemma (6 weeks pa) | Lodicule (6 weeks pa) | Palea (6 weeks pa) | Senescing leaf (2 months) |
| *Hv* CARa | 12.1096 | 13.1528 | 28.163 | 7.81317 | 21.6321 | 31.2757 | 40.3686 | 11.6433 | 15.858 | 13.4399 | 25.4358 | 7.30657 | 17.6147 | 7.04771 | 15.1406 | 25.776 |
| *Hv* CARb | 0 | 0.135488 | 0 | 0.21069 | 0 | 0.0361736 | 0.0408782 | 0 | 0 | 0 | 0.157723 | 0 | 21.7422 | 1.35837 | 32.544 | 0.204274 |
| *Hv* CARc | 0 | 0 | 0 | 0 | 0 | 0 | 0 | 0.0789594 | 0 | 0 | 0 | 0 | 4.07058 | 0 | 9.95602 | 0 |
| *Hv* CARd | 12.7542 | 9.14687 | 7.68059 | 10.7755 | 6.04315 | 13.7107 | 17.8717 | 16.8634 | 12.4718 | 11.1777 | 9.782 | 2.68701 | 13.1312 | 25.5659 | 19.7207 | 1.99491 |
| *Hv* CARe | 12.3452 | 20.7849 | 38.447 | 29.1917 | 25.5873 | 51.2172 | 31.7243 | 46.3371 | 9.56516 | 7.38431 | 17.6343 | 26.2672 | 46.9052 | 73.8073 | 31.5622 | 42.1943 |
| *Hv* CARf | 35.5278 | 21.7093 | 17.5014 | 15.0267 | 35.7123 | 38.2761 | 18.0313 | 13.4577 | 19.6631 | 8.5791 | 46.029 | 17.8842 | 10.6318 | 11.9449 | 8.24209 | 9.38378 |

**Table S4**: Primers used for cloning in this study

| Primer name | Primer sequence 5' -> 3' |
|---|---|
| attB1-CARa_F | GGGGACAAGTTTGTACAAAAAAGCAGGCTTAATGGAAGACGGGGCCG |
| attB1-CARa_R | GGGGACCACTTTGTACAAGAAAGCTGGGTTCACGGCCATCTCATTCTC |
| attB1-RIPb_F | GGGGACAAGTTTGTACAAAAAAGCAGGCTCAGGAATGCAGAACTCAAAAACCAGTAG |
| attB1-RIPb_R | GGGGACCACTTTGTACAAGAAAGCTGGGTCTCAGGTCTCATGAGCTTCTTCAC |
| attB1-RIPbCC2_F | GGGGACAAGTTTGTACAAAAAAGCAGGCTCAGAAATGCAGCCGGAGC |
| SpeI-CARa_F | AACTAGTAGATCCGTCCATCAGAGGATG |
| SalI-CARa_R | TGTCGACGGCGTCGGCTTCG |
| SpeI-RIPbCC2_F | AACTAGTTCCGAAATGCAGCCGGAGC |
| SalI-RIPbCC2_F | TCGGTCGGTCGACCGGTCTCATG |
| NotI-CARa-RNAi_F | GATATAGCGGCCGCGAGCGCAAAGACAAAAG |
| SalI-CARa-RNAi_R | GAGAGAGTCGACCTTGACGGTGATGAAGAG |
| SalI+CARa_F | GTCGACTTATGGAAGACGGGGCCGAG |
| NotI+CARa_R | GCGGCCGCCTACTTGGCGTCGGCTTCGATCCA |
| BamHI-RIPbCC2_F | GGATCCGAAATGCAGCCGGAGCTGGAGGC |
| EcoRI-RIPb_R | GGATCCATGCAGAACTCAAAAACCAGT |
